# Circulating glutamine and Alzheimer’s disease: a Mendelian randomization study

**DOI:** 10.1101/819029

**Authors:** Charleen D. Adams

## Abstract

**INTRODUCTION:** Alzheimer’s disease is a devastating neurodegenerative disorder. Its worldwide prevalence is over 24 million and is expected to double by 2040. Finding ways to prevent its cognitive decline is urgent.

**METHODS:** A two-sample Mendelian randomization study was performed instrumenting glutamine, which is abundant in blood, capable of crossing the blood-brain barrier, and involved in a metabolic cycle with glutamate in the brain.

**RESULTS:** The results reveal a protective effect of circulating glutamine (inverse-variance weighted method, odds ratio per 1-SD increase in circulating glutamine = 0.83; 95% CI 0.71, 0.97; *P* = 0.02).

**CONCLUSION:** These findings lend credence to the emerging story supporting the modifiability of glutamine/glutamate metabolism for the prevention of cognitive decline. More circulating glutamine might mean that more substrate is available during times of stress, acting as a neuroprotectant. Modifications to exogenous glutamine may be worth exploring in future efforts to prevent and/or treat Alzheimer’s disease.

## 1. Introduction

Recently, Zheng *et al*. (2019) reported a restoration of cognitive function in late-stage familial Alzheimer’s disease (FAD) mice with inhibition of euchromatin histone methyltransferases 1 and 2 (EHMT1/2) [1]. Doing so reversed histone hypermethylation (H3K9me2) at the promoters of the glutamate receptors and restored their expression. These are important findings that point to the targetability of EHMT1/2 in relation to glutamate receptor biology and, indirectly, possibly to the targetability of glutamate metabolism. Because the worldwide prevalence of Alzheimer’s disease is over 24 million and expected to double by 2040, with an aging global population [2], finding ways to prevent Alzheimer’s disease or reverse its cognitive decline is a crucial medical challenge of paramount public-health importance. To this effort, a two-sample Mendelian randomization (MR) study was performed to investigate the modifiability of an aspect of glutamate receptor biology—specifically, circulating glutamine—in the etiology of Alzheimer’s disease.

## 2. Methods

### 2.1. Conceptual approach

Briefly, MR is an instrumental variables technique that is analogous to a randomized controlled trial (RCT) and used to test mechanisms. Both RCTs and MR studies exploit random allocation to assess causal effects: for RCTs, this is done by random allocation of treatment by investigator, and for MR studies, random allocation occurs by ‘genetic lottery’ [3] – through the random assortment of alleles. MR can be done to investigate mechanisms due to a ubiquitous property of the human genome: pleiotropy – genetic variants influencing more than one trait. When a genetic variant influences a particular trait, which in turn directly influences another, this form of (vertical) pleiotropy can be exploited for causal inference with MR [4].

For the present analysis, in order to perform two-sample MR, some aspect of glutamate receptor biology needed to be instrumented. The glutamate receptors are difficult to genetically instrument, since few large-scale genome-wide association studies (GWAS) exist for measures of traits in tissues other than blood. However, downregulation of the glutamate receptor might impact the neurotransmitters which interact with the glutamate receptor, potentially implicating the glutamate/glutamine cycle. Since the glutamate/glutamine cycle is responsive to exogenous glutamine, which is abundant in blood and capable of crossing the blood-brain barrier [5,6], a GWAS of circulating glutamine can serve as the first data source for two-sample MR to instrument glutamine.

### 2.2. Data sources

*Step 1*. Kettunen *et al*. (2016) performed a GWAS of 123 circulating metabolites—including glutamine—in 24,925 participants from 14 European cohorts [7]. From this, independent (those not in linkage disequilibrium; R^2^ < 0.01) single-nucleotide polymorphisms (SNPs) associated at genome-wide significance (*P* < 5 × 10^−8^) with a standard-deviation (SD) increase in circulating glutamine were identified.

*Step 2*. A second GWAS data source—one for Alzheimer’s disease—the International Genomics of Alzheimer’s Project (IGAP) was selected [8]. The IGAP study contains 17,008 Alzheimer’s disease cases and 37,154 controls of European ancestry.

*Step 3*. Summary statistics (effect estimates, standard errors, and *p*-values) for the glutamine SNPs were extracted from the Kettunen GWAS. Four SNPs were available for this purpose. Likewise, summary statistics for these four SNPs were extracted from the IGAP GWAS (Table 1 contains characteristics of the SNPs).

**Table 1.**
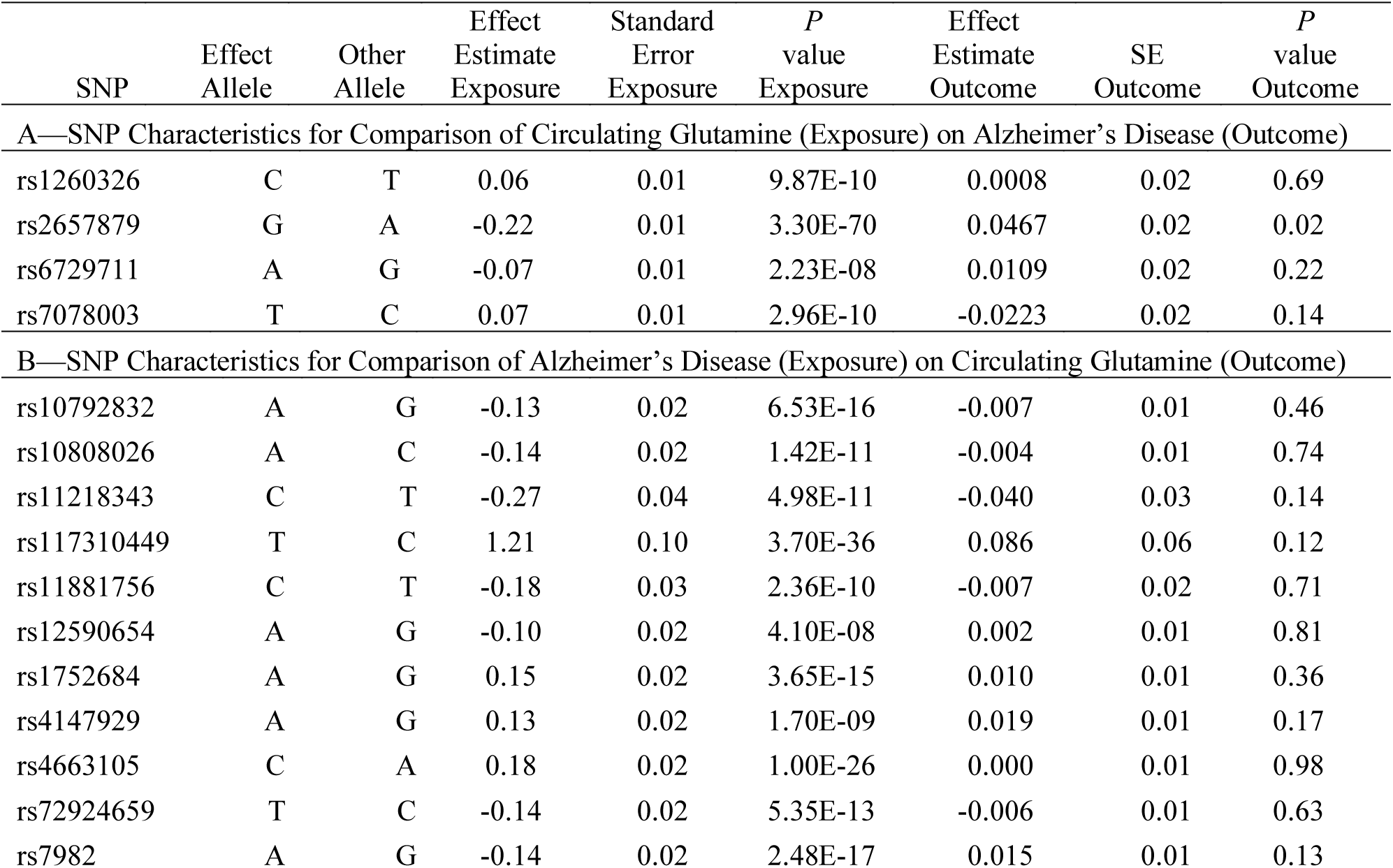
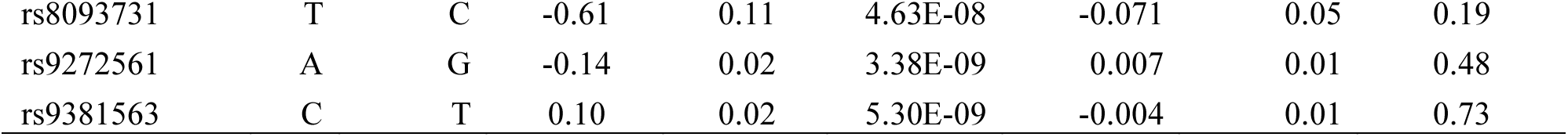
Two-sample Mendelian randomization instrument characteristics

### 2.3. Inverse-variance weighted (IVW) method

The summary statistics in Table 1 (panel A) were used to construct a four-SNP instrument for circulating glutamine. With this, the log-odds for Alzheimer’s disease per SD increase in circulating glutamine was calculated, using the inverse-variance weighted (IVW) MR method. With IVW, a weighted average is calculated using the inverse of the variance for the effects of the SNPs on Alzheimer’s disease as the weights. *X*_*z*_ is the effect of the SNP on glutamine, and *Y*_*z*_ is the effect of the SNP on Alzheimer’s disease. For the *z*^*th*^ variant, the IVW is defined as:

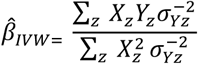

IVW treats each SNP as a natural experiment and provides a meta-analysis of the Wald ratios [9–12] (Fig. 1 describes the method in more detail). The results were exponentiated to obtain odds ratios (OR) for interpretability: OR<1 indicates a protective effect.

**Fig. 1.**
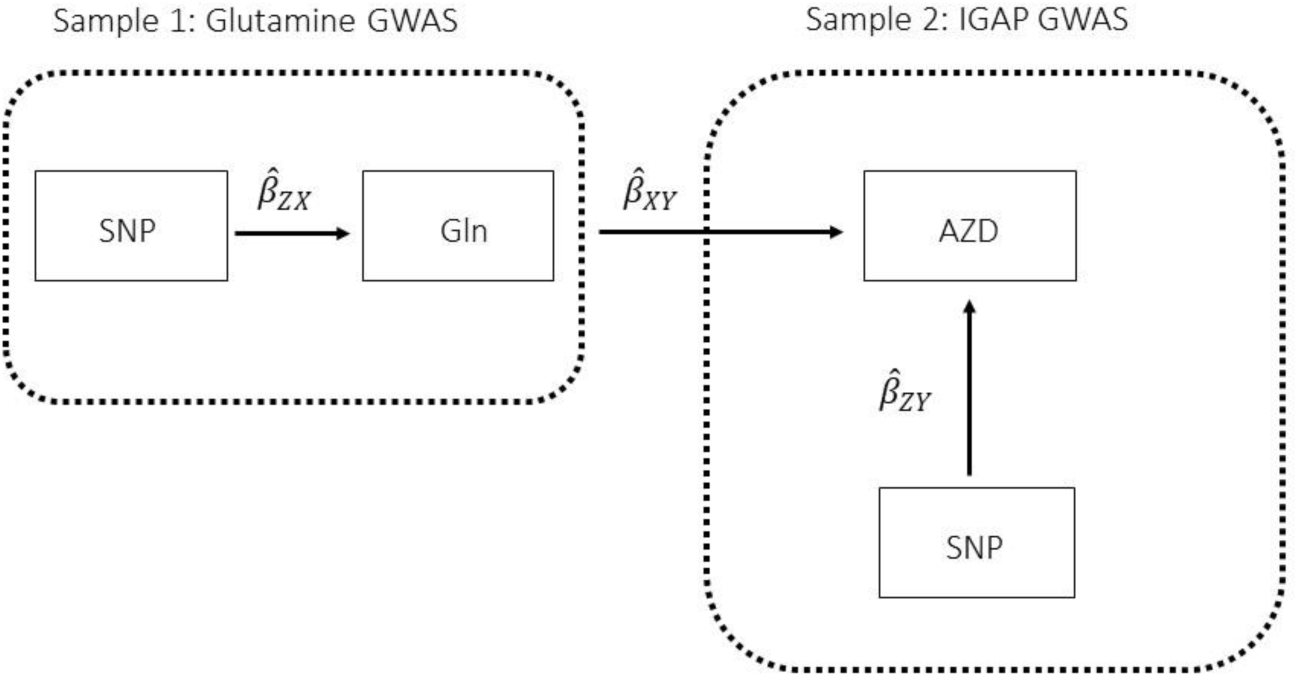
Two-sample Mendelian randomization testing the causal effect of circulating levels of glutamine (Gln) on Alzheimer’s disease (AZD). Estimates of the SNP-Gln association 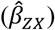 are calculated in sample 1 (Kettunen *et al*. (2016) GWAS). The association between these same SNPs and AZD 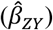 is then estimated in sample 2 (IGAP GWAS). These estimates are combined 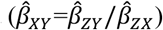. The 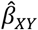 Wald ratio estimates for each of the four SNPs are meta-analyzed using the inverse-variance weighted 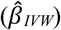 method and sensitivity analyses. The IVW method produces an overall causal estimate of circulating glutamine on AZD.

### 2.4. Sensitivity analyses

In order for the present MR results to be valid, three assumptions must hold: (i) the instrumental variable (IV) must be strongly associated with glutamine; (ii) the IV must be independent of confounders of glutamine and Alzheimer’s disease; and (iii) the IV must not be pleiotropically associated with Alzheimer’s disease; i.e., the IV must be associated with Alzheimer’s disease only through glutamine and not associated with Alzheimer’s disease through other exposures [13]. Assumption (iii) describes horizontal (a.k.a. “bad”) pleiotropy, which, when present, can invalidate the desired causal inference from vertical (a.k.a. “good”) pleiotropy. It potentially induces false-positive association and/or noise into the model, leading to a reduction in power to see a true effect [4].

As is standard, assumption (i) was dealt with by selecting SNPs strongly associated with circulating glutamine (genome-wide significance: *P* < 5 × 10^−8^). The use of SNPs as instruments greatly reduces the chance of confounding from environmental factors, addressing assumption (ii). To address assumption (iii), a sensitivity analysis was performed to screen for directional horizontal pleiotropy: MR Egger regression, weighted median, and weighted mode MR methods were run as complements to the IVW method. If the magnitudes and directions of the various MR methods comport across estimators, this lack of heterogeneity provides some evidence against pleiotropy (i.e., because the MR methods make different assumptions about the underlying nature of pleiotropy, it is unlikely that violations to the pleiotropy assumption would result in similar estimates across them all).

Briefly, the IVW method can be biased if any of the SNPs in its instrument suffer from horizontal pleiotropic effects, but MR Egger regression can provide an unbiased causal effect in the presence of such pleiotropy. Additionally, its intercept is not constrained to zero and can be used to estimate the average directional pleiotropic effect of the instrument [14]. Likewise, the weighted median estimator presumes 50% of the variants are invalid due to pleiotropy and provides a valid estimate in the face of this possibility [15]. Similarly, the weighted mode estimator can provide a robust causal estimate, even if the majority of instruments are invalid, when the largest number of similar causal effect estimates comes from valid instruments [16]. Supplementing these more widely used approaches, a robust adjusted profile score (RAPS), a recently developed MR method that is robust to idiosyncratic pleiotropy was run [17]. An extended description of the different MR methods and the different assumptions they make about pleiotropy is available elsewhere [14,18,19].

Lastly, beyond these sensitivity estimations to screen for violations to assumption (iii), further steps were taken. A leave-one-out permutation test was performed to assess whether the IVW estimate was biased by the influence of particular SNPs. Each of the four SNPs were likewise examined in PhenoScanner, a curated database of GWAS studies containing SNP-phenotype associations [20,21]. The SNPs were examined for associations with potential pleiotropic confounders.

### 2.5. Reverse direction

The hypothesized mechanism is an influence of glutamine on Alzheimer’s disease. However, because glutamine is capable of existing the brain as well as entering it [22], the reverse association (Alzheimer’s disease impacting glutamine levels) is also possible, making this a bi-directional MR study. Bi-directional MR requires that the instruments chosen for each tested direction are independent of each other; i.e., that there is no overlap in the SNPs for each instrument nor linkage disequilibrium between them [23,24]. To account this, LDassoc was used to verify independence [25].

The same GWAS data sources used for the test of glutamine on Alzheimer’s disease were used for the reverse association. Fourteen SNPs associated at genome-wide significance with Alzheimer’s disease were available in the IGAP GWAS and extractable in the Kettunen GWAS (SNP characteristics are in Table 1) for two-sample MR. The aforementioned IVW method and concomitant sensitivity analyses were performed. Due to performing a bi-directional association (two tests), a Bonferroni correction was used to set the significance threshold (0.05/2 = 0.025) for the IVW associations.

### 2.6. Power and MR tools

Power for the analysis was based on the primary hypothesis: glutamine on Alzheimer’s disease. The proportion of the variance in glutamine explained by the four-SNP instrument (R^2^) and the strength of the instrument (*F* statistic) were calculated. For the *F* statistic, the following formula was used, where *n* indicates the sample size and *k* denotes the number of SNPs:

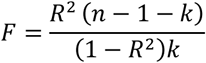

*F* statistics <10 indicate that the instrument is weak. Weak instruments can bias the findings[26,27].

The MR study was powered using the mRnd MR power calculator (available at https://cnsgenomics.shinyapps.io/mRnd/) [28] and the MR analysis was run in R version 3.5.2 using the TwoSampleMR [29] and MendelianRandomization [30] packages.

## 3. Results

### 3.1. Glutamine on Alzheimer’s disease

The results reveal a protective effect of circulating glutamine on Alzheimer’s disease (OR per 1-SD increase in circulating glutamine: IVW estimate 0.83; 95% CI 0.71, 0.97; *P* = 0.02; Fig. 2). The R^2^ for the glutamine instrument was 0.019 and the *F* statistic was 119.

**Fig. 2.**
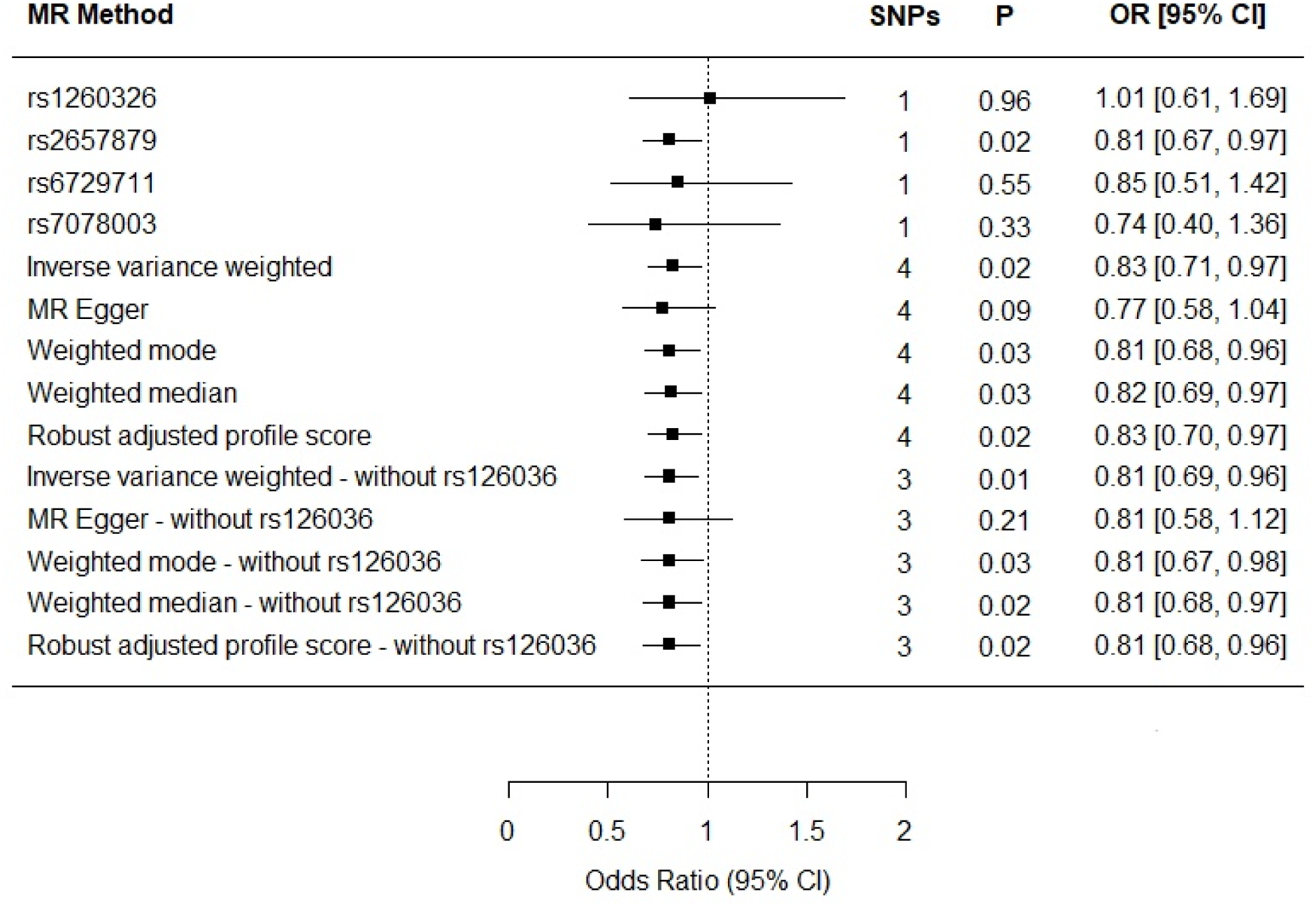
Individual SNP and multi-instrument MR results for the effect of circulating glutamine on Alzheimer’s disease. In addition to the test of the causal effect of glutamine on Alzheimer’s disease (depicted above), the MR Egger method provides a test of whether its intercept is different from zero. MR Egger intercept = 0.01; 95% CI −0.03, 0.04; *P* value=0.60 (four-SNP instrument).

### 3.2. Alzheimer’s disease on glutamine

The results for the reverse association, Alzheimer’s disease on glutamine, were null (IVW β estimate for liability to Alzheimer’s disease = 0.03; 95% CI −0.01, 0.07; *P* = 0.12; Fig. 3).

**Fig. 3.**
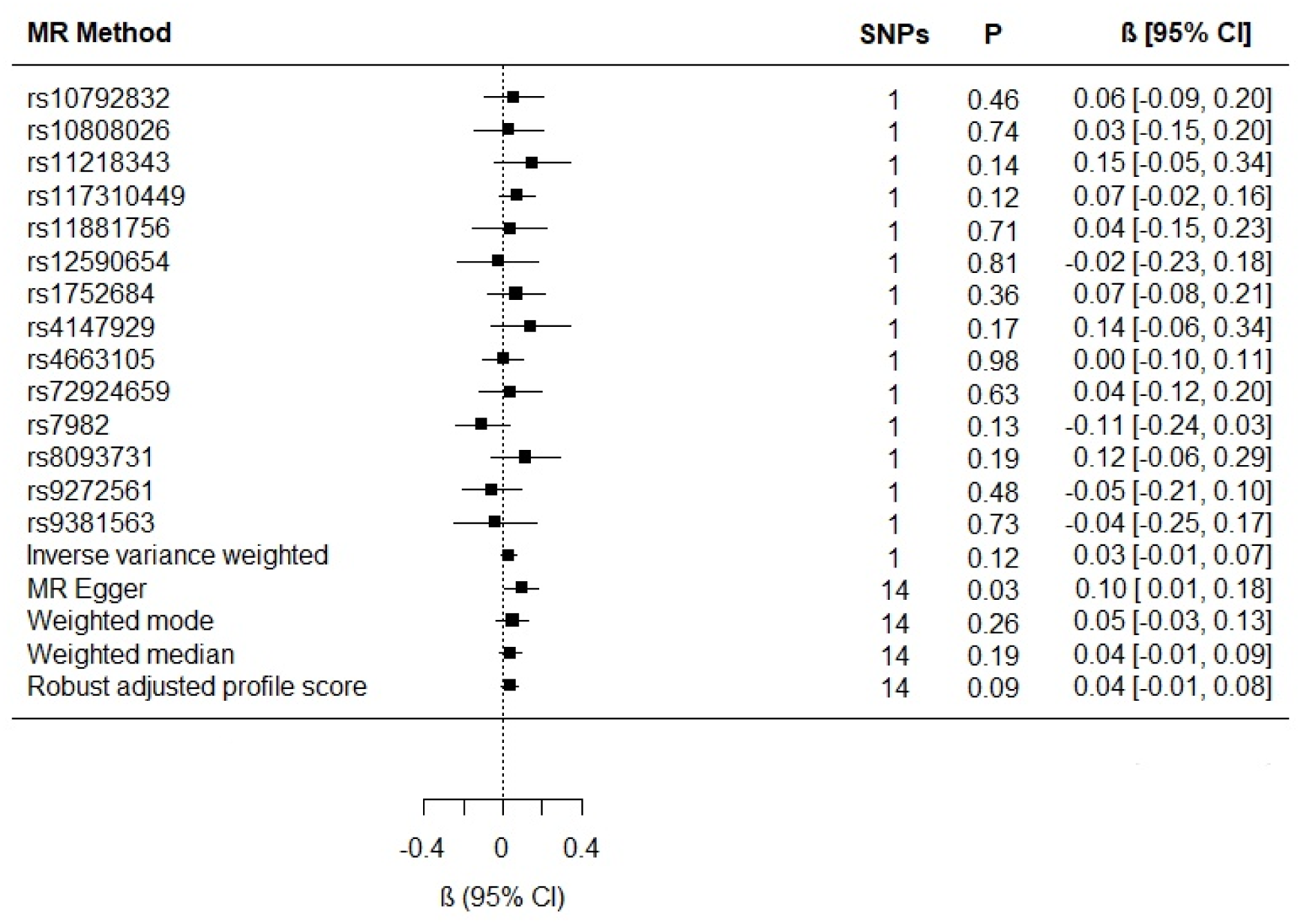
Individual SNP and multi-instrument MR results for the effect of Alzheimer’s disease on circulating glutamine. MR Egger intercept (−0.01; 95% CI −0.03, 0.002; *P* value = 0.09).

## 4. Discussion

### 4.1 Strengths and limitations

With the proportion of cases in the IGAP GWAS being 31% and an alpha threshold of 0.05, the study was aptly powered for the study of glutamine on Alzheimer’s disease; there was 89% power to detect a true effect of 0.80, pointing to a strength of this analysis: use of large GWAS data sources made it possible to capitalize on the power of large samples to detect effects. Moreover, because of the two-sample MR design, potential bias from weak instruments would have bent the results towards the null [26], reducing concerns about false-positive findings.

A general limitation of MR is the persistent possibility of horizontal pleiotropic association between an instrument and an outcome, independent of the exposure of interest – in this case, the possibility that the glutamine SNPs impact Alzheimer’s disease through an intermediate phenotype other than glutamine. However, an attempt to separate out the vertical pleiotropic (i.e., hypothesized) pathway from these other influences was made. The complementary sensitivity estimators left the causal association between glutamine and Alzheimer’s disease essentially unchanged: notwithstanding a slightly more protective effect with the MR Egger estimate (0.77 vs 0.83), the weighted median and weighted mode estimations were consistent with those of the IVW in terms of direction and magnitude of effects, and there was confirmation of the IVW findings from the RAPS estimator (Table 1).

The MR Egger estimate lacked precision, but MR Egger’s power to detect causal effects is less than IVW’s [11]. In addition, it’s lack of precision does not contradict the evidence for causality obtained from the IVW method [31]. The *p*-value for the MR Egger intercept test provided evidence against bias in the IVW estimate (MR Egger intercept estimate = 0.01; 95% CI −0.03, 0.04; *P* =0.60). Cochran’s *Q*-statistic (a formal test of heterogeneity in SNPs) was rejected, implying no pleiotropy (heterogeneity estimate = 0.53; *P* = 0.77). The 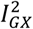 statistic (97.2%), a measure of instrument strength specifically for two-sample MR, indicated relatively little bias towards the null in the MR Egger causal estimate [14,32].

The leave-one-out permutation analysis provided no evidence that the results were driven by particular SNPs (OR 0.83; 95% CI 0.71, 0.97; *P* = 0.02). Nonetheless, the PhenoScanner results revealed that rs1260326 was also associated with Apolipoprotein A1 (APOA1), a known risk factor for Alzheimer’s disease. To determine whether removing this pleiotropic effect impacted the results, the analysis was re-run removing rs1260326. Doing so increased the strength of the protective effect in comparison to the IVW estimate including rs1260326 (OR = 0.83): rs1260326removed IVW OR 0.81; 95% CI 0.69, 0.96; *P* = 0.01; Fig. 2) and resolved the remaining discordance between the IVW and MR Egger estimates: all the estimators with rs1260326 removed produced an effect size of 0.81.

Overall, the various sensitivity analyses demonstrated little evidence of horizontal directional pleiotropy biasing the observed protective effect of glutamine on Alzheimer’s disease, especially after rs1260326 was removed.

There was general consistency between the sensitivity estimators in terms of the magnitude and direction of the effects, but a discordance between the IVW and MR Egger tests: the MR Egger estimate revealed a causal association (MR Egger β = 0.10; 95% CI 0.01, 0.18; *P* = 0.03; Fig. 3). However, because MR Egger is a sensitivity test and not designed to stand on its own, the primary null association observed with the IVW method (β = 0.03; 95% CI −0.01, 0.07; *P* = 0.12; Fig. 3) is the conventional metric for appraising causality [31]. While there was no evidence of directional pleiotropy from the MR intercept test (MR Egger intercept estimate = −0.01; 95% CI −0.03, 0.002; *P* = 0.09) and Cochran’s *Q*-statistic was rejected (heterogeneity estimate = 7.82; *P* = 0.80), it is possible for the MR Egger method to provide false-positive results, if the average pleiotropic effects are not independent of the SNP associations with Alzheimer’s disease – i.e., if there is a violation to the InSIDE assumption (INstrument Strength Independent of Direct Effect) [31,33]. Additionally, the 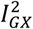 statistic (77.5%) indicated that the instrument may have been underpowered: 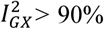, or a bias towards the null of 10%, is the conventional threshold for determining the appropriate instrument strength for MR Egger [32]. Due to this, a causal association for the effect of Alzheimer’s disease on glutamine could not be ruled in.

### 4.2. Biological Plausibility

A recent cohort study of 1356 Alzheimer’s disease patients and 23,882 controls found an increased risk for Alzheimer’s disease correlating with higher circulating levels of glutamine (OR 1.11; 95% CI 1.04–1.20; *P* = 0.003). However, observational studies are prone to reverse causation and confounding. It is possible that their findings reflect the egress of glutamine from the brain across the blood-brain barrier; hence, they may have observed a consequence, not a cause, of the disease.

Similarly, a recent case-control study observed an increase in cerebral spinal fluid (CSF) glutamine in 72 probable Alzheimer’s disease cases (all scoring positive on the amyloid tau index, a biomarker of amyloid-β and tau neuropathology) versus 71 age-matched controls. Their data suggest that CSF glutamate and glutamine levels might serve as a subclinical biomarker of subtle cognitive changes [34]. Further to this, in the brain, most endogenous glutamine is produced by glutamine synthetase (GS). Increased levels—but compromised activity—of GS may explain the increased levels of CSF glutamine in Alzheimer’s disease patients [5].

Notwithstanding the possibility that circulating (and CSF) levels of glutamine may reflect a consequence of Alzheimer’s disease, it is biologically plausible for glutamine to act mechanistically in the etiology of the disorder. Chen and Herrup (2012) found that 1) raising glutamine levels in cultured cells protected them from amyloid peptide; 2) providing glutamine supplementation to two animal models of Alzheimer’s disease decreased biochemical indices of dysfunction (namely, inflammation-induced neuronal cell cycle activation, tau phosphorylation, and ATM-activation); and 3) glutamine deprivation decreased autophagy [5]. Their results imply that exogenous glutamine plays a protective role in neuronal health. Moreover, Anderson *et al*. (2017) observed that reduced glutamine uptake and hampered oxidative glutamine metabolism precede amyloid plaque formation in APPswe/PSEN1dE9 mice compared to controls. This implies that alterations in glutamine metabolism may be early transformations in the pathogenesis of Alzheimer’s disease [35]. Since the accumulation of amyloid plaques has been associated with inflammation, oxidative stress and mitochondrial dysfunction [36], perhaps exogenous glutamine can protect against this in the early phrases of the disease.

## 5. Conclusion

The present findings lend credence to the emerging story supporting the modifiability of key aspects of glutamate/glutamine metabolism, and, indirectly, the observations by Zheng *et al*. (2019), who found that restoration of the expression of the glutamate receptor (via EHMT1/2 inhibition) reversed cognitive deficits in their FAD mice [1]. A higher circulating level of glutamine might mean that more of the substrate is available for use in the brain during times of stress, acting as a neuroprotectant [5]. Modifications to glutamine may be worth exploring in future efforts to prevent this devastating disease.

## Acknowledgements

The author thanks Kettunen *et al.* (2016) GWAS and the IGAP consortium for making their data public.

